# Evolution of molecular determinants for SUMO-activating enzyme subcellular localization in plants

**DOI:** 10.1101/2020.10.05.326249

**Authors:** Abraham Más, Laura Castaño-Miquel, Lorenzo Carretero-Paulet, Núria Colomé, Francesc Canals, L. Maria Lois

**Affiliations:** Centre for Research in Agricultural Genomics (CRAG) CSIC-IRTA-UAB-UB, Campus UAB, Bellaterra, Barcelona, Spain; Consejo Superior de Investigaciones Científicas (CSIC), Barcelona, Spain; Department of Biology and Geology. University of Almería. Ctra. Sacramento s/n 04120 Almería, Spain; Vall d’Hebron Institute of Oncology (VHIO), Barcelona, Spain

**Keywords:** SUMO E1-activating enzyme, subcellular localization, seed evolution, proteolytic processing

## Abstract

Post-translational modification by Small Ubiquitin-related Modifier (SUMO) is an essential regulatory mechanism in eukaryotes. In the cell, SUMO conjugates are highly enriched in the nucleus and, consistently, SUMOylation machinery components are mainly nuclear. Nonetheless, cytosolic SUMO targets also exist and the mechanisms that facilitate SUMO conjugation in the cytosol are unknown. Here, we show that the nuclear localization of the *Arabidopsis* SUMO activating enzyme large subunit SAE2 is dependent on two nuclear localization signals, the canonical NLS1 and the non-canonical NLS2 identified and validated here. NLS2 is proteolytic processed from SAE2 during seed development, facilitating SAE2 enrichment in the cytosol. Results obtained using transgenic plants expressing different SAE2 proteoforms suggest that SAE2 cytosolic enrichment could constitute a rapid signal for growth arrest. Phylogenetic studies indicated that the *Arabidopsis* NLS1-NLS2 structural organization is conserved only in seed plants, providing a potential evolutionary role of cytosolic SUMOylation in seed appearance.

## Introduction

Among the multilayer mechanisms that mediate signal integration to environmental and endogenous cues in eukaryotes, post-translational modifications (PTMs) facilitate a highly dynamic and rapid response^1^. PTMs consist in the reversible addition of small molecules or proteins to specific residues in the target proteins, or in the irreversible limited proteolysis occurring at precise peptide linkages^2^. PTMs provide an increased complexity and precision to cellular responses by modifying protein function. Protein modification by SUMO (small ubiquitin-related modifier), or SUMOylation, consists in the addition of a small protein to a lysine residue in the target protein. In plants and animals, SUMOylation plays an essential role during embryo development^3,4^. The seed developmental stage provide plants with the capacity to survive to adverse environmental conditions and the colonization of new habitats. Seed development starts with embryogenesis and embryo growth, followed by the seed maturation phase, which is characterized by embryo growth arrest, accumulation of food reserves and development of dormancy^5^. Beyond early stages of embryogenesis, SUMO has a major role in plant development and tolerance to biotic and abiotic stress^6–9^. The fact that SUMOylation regulates many plant traits of agronomic interest has stimulated the generation of multiple approaches aiming to modulate plant SUMOylation as strategy to increase crop productivity^10^.

SUMO is conjugated as a monomer at one single site or multiple sites, and/or forming polySUMO chains that are further modified by SUMO-targeted Ubiquitin ligases^11^, which provides a mechanism for proteolytic control of SUMOylated proteins. SUMOylation is a highly regulated process mediated by the sequential action of the E1-activating enzyme, E2-conjugating enzyme and an E3 ligase (Figure 1a). The heterodimeric E1-activating enzyme catalyzes the ATP-dependent first step in SUMO conjugation. The E1 consists of a small subunit, SAE1, and a large subunit, SAE2, which is structured in four functional domains: adenylation, catalytic cysteine, ubiquitin fold (UFD) and the C-terminal domains^12^ (Figure 1b). As reversible modification, SUMO is removed from the substrates by specific proteases, ULPs^13^ (Figure 1a).

**Figure 1.**
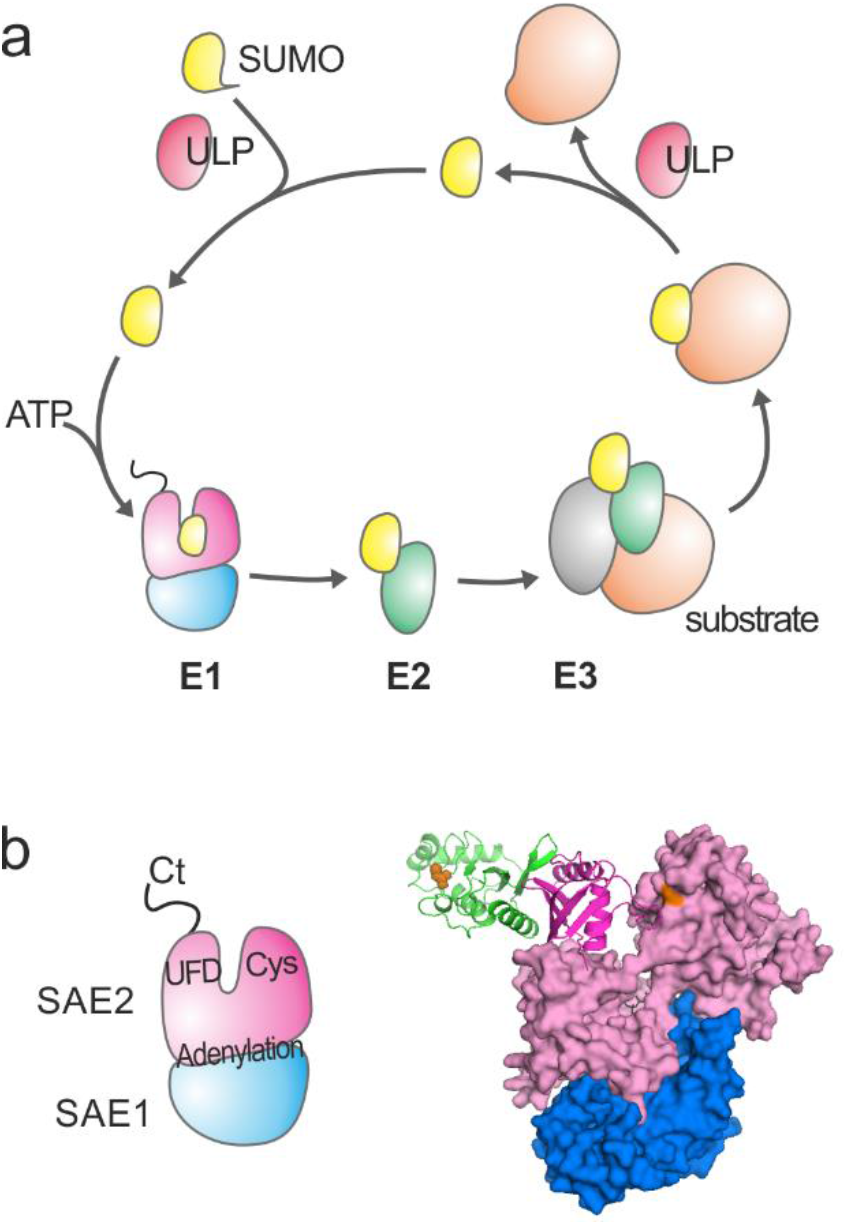
SUMO conjugation/deconjugation machinery components. (a) SUMO conjugation/deconjugation cycle. SUMO is synthesized as a precursor that is processed at its C-terminal tail by the specific ULP proteases, releasing a SUMO mature form with a Gly–Gly motif at its C-terminus. Subsequently, SUMO is activated by the heterodimeric E1 activating enzyme, SAE1/SAE2, transferred to the E2 conjugating enzyme, and, finally, attached to a target lysine in the substrate. The target lysine is usually located within the consensus site ΨKxE/D (Ψ is a large hydrophobic amino acid, and x, any amino acid). This final step is facilitated by E3 ligase enzymes that interact with both SUMO-charged E2 and the substrate. SUMOylation is a reversible modification, and the same class of cysteine proteases involved in the maturation step catalyze SUMO excision from the substrate. (b) Cartoon representation of the SUMO1 structural and functional organization (left). Model of *Arabidopsis* SUMO E1-E2 complex generated by alignment of molecules with the PDB IDs 1Y8Q and 6GUM using the SWISS-MODEL comparative protein modeling server. Adenylation and catalytic cysteine domains are shown in Gaussian surface representation. UFD and its interacting partner, SUMO E2, are in cartoon representation. SAE2 C-terminal tail is absent in the electron density maps of the PDBs available of the SUMO E1s.

In *Arabidopsis*, the E1 regulates SUMO conjugation by selecting SUMO isoforms entering to the conjugation pathway^14^ and, the conjugation rate by evolutionary diversification of SAE1 gene family, encoding for the E1 small subunit^15^. Similarly to the human and yeast orthologs, the *Arabidopsis* E1 displays nuclear localization facilitated by a signal located at the C-terminal tail^15^. Other components of the *Arabidopsis* SUMOylation machinery display distinct subcellular distributions^6,16^, although the biological relevance of their subcellular compartmentalization is unknown. In addition, different molecular mechanisms, ranging from gene diversification^14,15,17^ to post-translational modifications of the machinery components^18–23^, account for the complexity of the SUMOylation system in plants. More efforts aiming to understand the complex molecular circuit driving *in vivo* SUMOylation will uncover critical aspects required for precise engineering of SUMOylation to improve plant performance.

In this study, we have identified a novel post-translational mechanism targeting SUMO-activating enzyme large subunit, SAE2, in *Arabidopsis*. We have determined that the C-terminal eight amino acids of SAE2 undergo proteolytic cleavage during seed development, being concomitant to the stage of embryo growth arrest and the transition to seed maturation. The functional characterization of the released eight amino acids indicated that they constitute a novel non-canonical NLS2 that is essential, together with the canonical NLS1, for complete nuclear localization. The study of *Arabidopsis* transgenic plants expressing SAE2 proteoforms displaying different degrees of cytosolic localization suggests that SAE2 enrichment in the cytosol could constitute a growth arrest signal. We postulate that this regulated nucleo-cytosolic E1 shuttling modulates subcellular SUMO conjugation *in vivo* to coordinate plant responses to endogenous and exogenous stimuli. We found that the structural organization of the SAE2 C-terminal tail is distinct between seed and non-seed plants. While seed plants feature a long and poorly conserved linker region between NLS1 and NLS2, non-seed plants display a short stretch of 5-10 amino acids that separates both NLSs. Phylogenetic analysis supports that the emergence/acquisition of the SAE2 C-terminal tail structural signature (linker region) is concomitant to seed appearance in evolution, suggesting that SAE2 processing and cytosolic SUMOylation could have a major role in seed development.

## Results

### Post-translational processing of the SUMO activating enzyme large subunit, SAE2, C-terminus

During the characterization of plants with impaired SUMOylation capacity by expression of the SAE2 ubiquitin fold domain^24^, SAE2^UFDCt^, we observed that both endogenous SAE2 and recombinant SAE2^UFDCt^ displayed distinct electrophoretic mobility when comparing protein extracts from green siliques or rosette leaves. This tissue-dependent SAE2 differential mobility was also observed in Col0 plants (Figure 2b), pointing to the existence of an unknown post-translational modification that targets the UFDCt domain of SAE2. The SAE2^UFDCt^ domain plays an important role in E1 activity, being essential for establishing non-covalent interactions with the E2 through the UFD domain (Figure 1b) and determining nuclear localization through the C-terminal tail ^12,15^. To elucidate the molecular basis of such mobility shift, either targeted proteolytic processing or other post-translational modifications, recombinant His:SAE2^UFDCt^ protein was incubated in the presence of silique or rosette protein extracts from Col0 plants. Figure 2c shows that, in presence of silique protein extracts, recombinant SAE2^UFDCt^ mobility shifted into a lower molecular weight form in a time-dependent manner, pointing to the existence of a silique-specific SAE2^UFDCt^ proteolytic cleavage.

**Figura 2.**
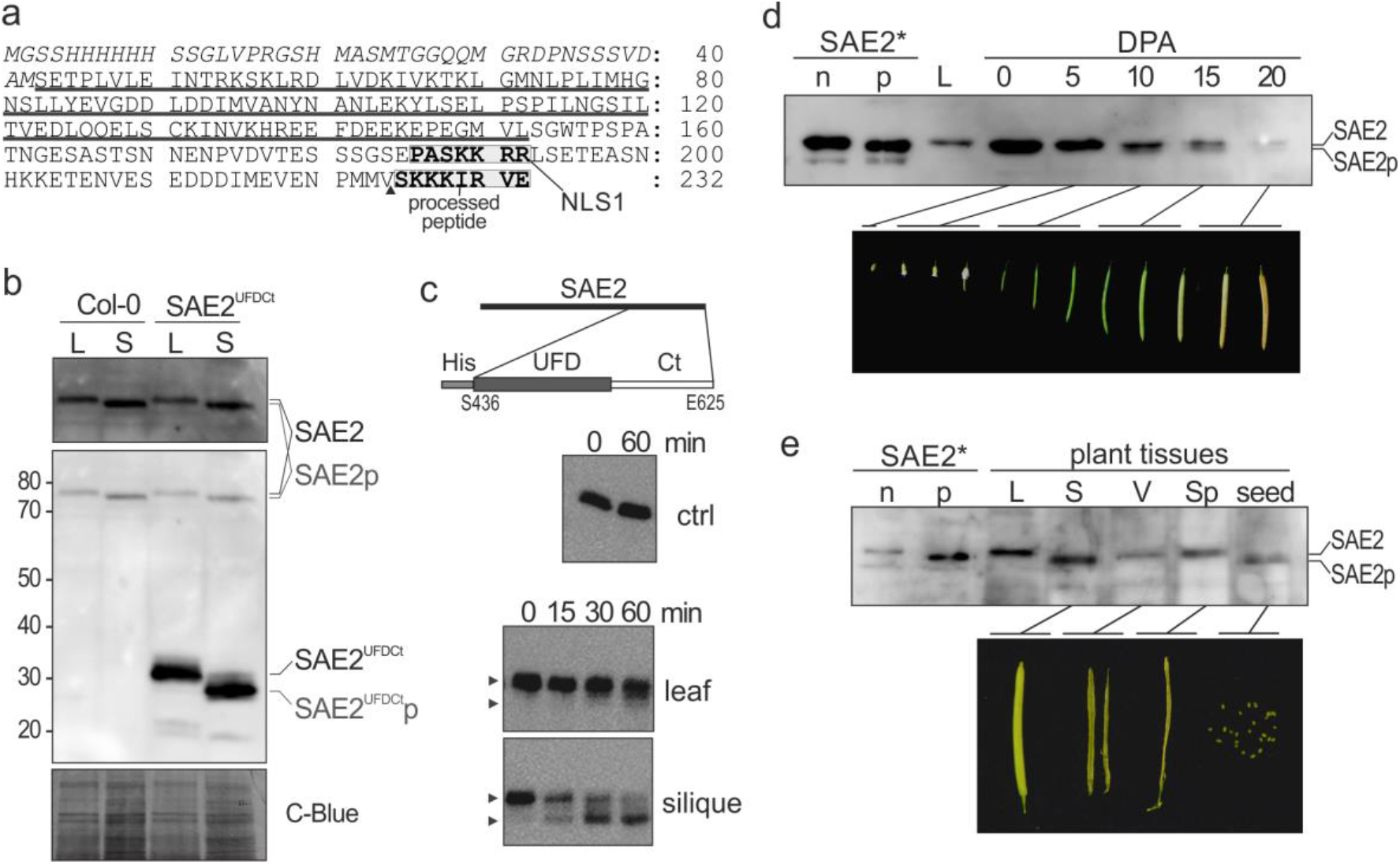
SAE2 C-terminal post-translational modification occurs in a tissue-dependent manner. (a) His:SAE2^UFDCt^ amino acid sequence corresponding to the recombinant protein used in C. The histidine tag is in italics. The SAE2^UFD^ domain is underlined. The amino acids corresponding to the canonical nuclear localization signal (NLS1) and the released amino acids after processing are enclosed in boxes. (b) SAE2 processing *in planta* identification in total protein extracts from rosette leaves (L) and siliques (S) from 5-week old Col-0 and SAE2^UFDCt^ expressing plants. (c) In vitro reconstitution of SAE2 proteolytic processing. 10 μg of recombinant His:SAE2^UFDCt^ were incubated at 30°C in the presence of 350 μg rosette and silique protein extracts from Col-0 plants. At the specified time points, aliquots were removed and denatured in the presence of loading buffer. (d) Temporal analysis of SAE2 proteoforms in seed development. Total protein extracts were prepared from Col0 flowers and siliques collected at different developmental stages as indicated. (e) Analysis of SAE2 proteoforms in dissected siliques. Total protein extracts were prepared from Col0 intact mature green siliques (S), and dissected valves (V), septum (Sp) and seeds. As controls, protein extracts from 7-week old rosette leave (L), and recombinant non-processed (n) and processed (p) SAE2 were included in (d) and (e). Protein extracts were resolved in 10% SDS-PAGE and SAE2 detected by immunoblotting using anti-SAE2 antibodies (b, c, d, e).

In order to identify the processing site, we performed preparative scale reactions and processed and unprocessed SAE2^UFDCt^ products were analyzed by mass spectrometry. The obtained spectra identified the peptide linkage between Val 617 and Ser 618 as the SAE2^UFDCt^ proteolytic site, releasing the SAE2 C-terminal eight amino acids (Figure 2a). In order to validate the processing site, recombinant protein of the two proteoforms SAE2 (Met1-E625) and processed SAE2 (Met1-V617) were produced in *E. coli* and their mobility on SDS-PAGE was compared with endogenous SAE2 present in silique and rosette leaves protein extracts. The results obtained by immunoblot analysis with anti-SAE2 antibodies were consistent with the identified processing site (Supplementary Fig. 2).

A detailed analysis of SAE2 proteoforms dynamics during seed development showed that SAE2 processing is observed in 10-day post-anthesis (DAP) and older siliques, suggesting that it could have an important role in the transition between embryo development and seed maturation. The dissection of 10-15 DAPs siliques showed that the SAE2 processed form is the most abundant variant in seeds, while in the other fruit structures, valves and septum, only the non-processed form is observed (Figure 2e).

### SAE2 limited proteolytic processing releases a non-canonical NLS required for complete nuclear localization

The enrichment in basic amino acids of the SAE2 processed sequence suggested that it could have functions of nuclear localization signal (Figure 2a). Previous studies showed that the heterodimeric SUMO E1-activating enzyme localizes preferentially to the nucleus and that this localization was facilitated by the C-terminal tail of the SAE2 subunit^15^, which contains a canonical NLS as identified by bioinformatic analyses. In mammals, similar studies showed that a nucleo-plasmin type bipartite NLS located in E1 large subunit was crucial for E1 nuclear targeting^25^. The putative consensus sequence corresponding to a bipartite NLS consists of two basic amino acid clusters separated by 10-12 residues that bind simultaneously to two binding pockets in importin α^26^. In *Arabidopsis* SAE2, 32 amino acids separate the conserved NLS from the eight C-terminal processed amino acids, suggesting the presence of two independent NLS: a canonical NLS and non-canonical NLS, which we have designated as NLS1 and NLS2, respectively. To test this hypothesis, we analyzed the subcellular localization of specific SAE2 domains transiently expressed as EYFP (enhanced yellow fluorescent protein) fusion proteins in onion cells. Analysis of relative cytosolic enrichment determined that the SAE2^NLS1^ peptide, the canonical NLS, confers nuclear enrichment to the EYFP, although not as efficiently as the SAE2^Ct^ tail or the SAE2^UFDCt^ domain. On the other hand, the SAE2^NLS2^ peptide is also competent to confer significant nuclear enrichment, although to a lesser extent than the SAE2^NLS1^ peptide (Figure 3a). Equivalent EYFP protein fusions were produced in *E. coli* to analyze their capacity to interact with native α-importin and mutant variants from yeast. In native gel-shift assays, both NLS1 and NLS2 peptides, as well as the SAE2 C-terminal tail, interacted mainly with the major groove of yeast α-importin, suggesting that they do not establish simultaneous contacts with different pockets of the α-importin during nuclear translocation, and that they do not behave as a bipartite NLS (Figure 3b).

**Figure 3.**
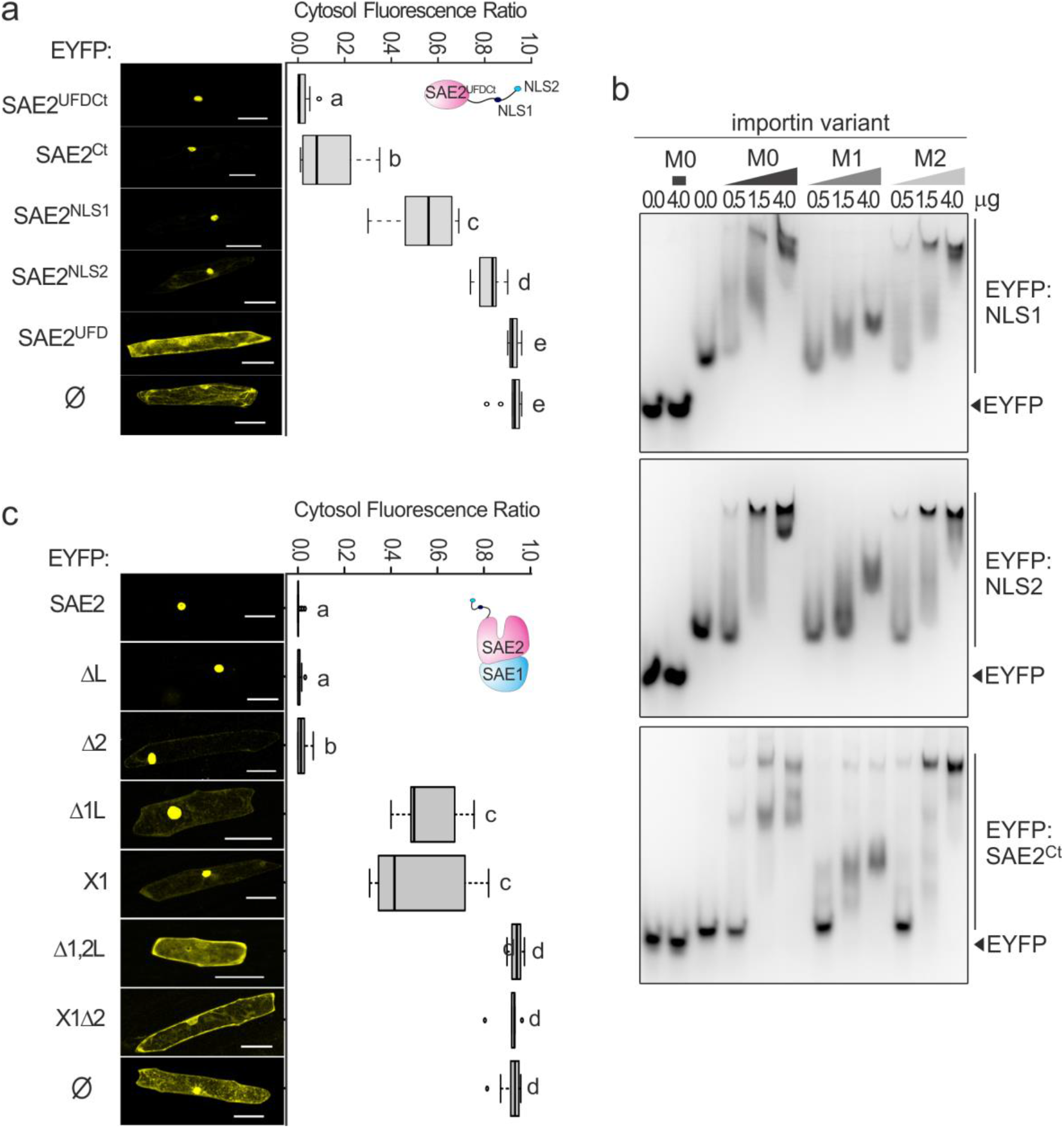
SAE2 nuclear localization is mediated by two nuclear localization signals, NLS1 and NLS2, displaying equivalent importin binding properties. (a) SAE2^UFDCt^ domain or the indicated variants were fused to EYFP and transiently expressed in onion epidermal cells. NLS1 sequence corresponds to P_579_ASKKRR and NLS2 sequence corresponds to S_618_KKKIRE. EYFP was used as control (Ø). Fluorescence from cytosol and nucleus was quantified and the cytosolic enrichment ratio calculated. Obtained values were plotted on the graph. A minimum of 8 cells were analyzed for each protein variant. Center lines show the medians; box limits indicate the 25th and 75th percentiles as determined by R software; whiskers extend 1.5 times the interquartile range from the 25th and 75th percentiles, outliers are represented by dots. T-test was performed and groups with different letters denote statistically significant differences between them (T-test *P* < 0.02). Bars= 100 μm. (b) Full length SAE2 or the indicated variants were fused to EYFP and transiently expressed in onion epidermal cells. EYFP was used as control (Ø). Fluorescence from cytosol and nucleus was quantified and the cytosolic enrichment ratio calculated. Obtained values were plotted on the graph. The number of quantified cells were as follow: 10 expressing EYFP (*Ø*); 17 expressing EYFP:SAE2; 8 expressing EYFP:SAE2ΔL; 18 expressing EYFP:SAE2Δ2; 8 expressing EYFP:SAE2Δ1L; 7 expressing EYFP:SAE2X1; 7 expressing EYFP:SAE2Δ1,2L; and 6 expressing EYFP:SAE2X1Δ2. Letters denote statistically significant differences between samples (*P* < 0.03). Bars= 100 μM. (c) Increasing amounts of native α-importin (Kap60pΔIBB, M0) from yeast or its mutant variants defective in NLS binding through the major (Kap60pΔIBB W195A/N199A, M1) or minor grooves (Kap60pΔIBB W405A/N409A, M2) were incubated in presence of 1 μM of His:2xflag:EYFP or His:2xflag:EYFP fused to SAE2^NLS1^ (top), SAE2^NLS2^ (middle) or SAE2^Ct^ (bottom). Complex formation was analyzed by native protein electrophoresis and detected by EYFP fluorescence emission under UV. Assays containing EYFP were analyzed in all protein gels as negative control. A representative image from 3 independent assays is shown.

To further evaluate the role of the SAE2^NLS2^ peptide as a *bona fide* nuclear localization signal, we analyzed its contribution to the E1 holoenzyme nuclear localization. Full-length SAE2 or SAE2 variants with single, Δnls1 or Δnls2, or double, Δnls1-2, NLSs deletions were fused to the EYFP C-terminus and their localization analyzed in transient expression experiments in epidermal onion cells. Consistently with previous results, full length SAE2 also displayed an exclusive nuclear localization. In contrast, significant fluorescence signal was detected in the cytosol when EYFP:SAE2Δnls1 or EYFP:SAE2Δnls2 were expressed, although the ratio nuclear:cytosolic distribution was dependent on the retained NLS. EYFP:SAE2Δnls1 displayed a stronger cytosolic localization than EYFP:SAE2Δnls2, suggesting that NLS1 contribution to E1 nuclear localization is quantitatively more relevant than NLS2. Moreover, the cytosolic localization of EYFP:SAE2Δnls1-2 suggests that NLS1 and NLS2 are the main determinants for SAE2 nuclear localization and that both signals are necessary to recapitulate SAE2 nuclear localization (Figure 3b). Equivalent results were obtained when we analyzed SAE2 NLS1 mutated variants (K582T K583G) alone or in combination with NLS2 deletions. The essential role of the NLS2 sequence for complete nuclear localization was also evident for the SAE2^UFDCt^ domain (Supplementary Fig. 3). Supporting our hypothesis that suggests a SAE2 nucleo-cytosolic shuttling, non-targeted proteomic studies identified SAE2 in the cytosolic protein fraction from *Arabidopsis* cell suspensions^27^.

### Biological significance of SAE2 C-terminal tail proteolytic processing

For uncovering the biological role of SAE2 proteolytic processing, we characterized *Arabidopsis* transgenic lines expressing SAE2 proteoforms differentially enriched in the cytosol (Figure 4 a, c), under the control of the endogenous SAE2 promoter (EYFP:SAE2, EYFP:SAE2X1, EYFP:SAE2Δ2 and EYFP:SAE2X1Δ2) or the 35S promoter (EYFP:SAE2, EYFP:SAE2Δ1L, EYFP:SAE2Δ2, EYFP:SAE2Δ1,2L) (Supplementary Figure 4). Two independent homozygous transgenic lines for each SAE2 proteoform were generated. As selection criteria, we aimed to identify independent lines displaying SAE2 proteoform levels as similar as technically possible. SAE2, SCE1 and free SUMO and SUMO conjugates levels were analyzed in dissected roots and shoots from 11-day plants. No significant differences were observed in SCE1 levels in any of the analyzed sample, (Figure 4a and Supplementary Figure 5). As expected, SAE2 proteoform levels over-accumulated in comparison to endogenous SAE2 when expressed under the control of the 35S promoter, with the exception of the fully cytosolic EYFP:SAE2Δ1,2L proteoform that accumulated to a much lower level (Supplementary Figure 5). These results suggest the existence of a molecular surveillance mechanism to maintain low SAE2 levels in the cytosol. In plants expressing SAE2 proteoforms under the SAE2 promoter, these differences in protein accumulation between nuclear and cytosolic SAE2 forms were not evident, probably as a result of their lower expression levels (Figure 4 a, b).

**Figure 4.**
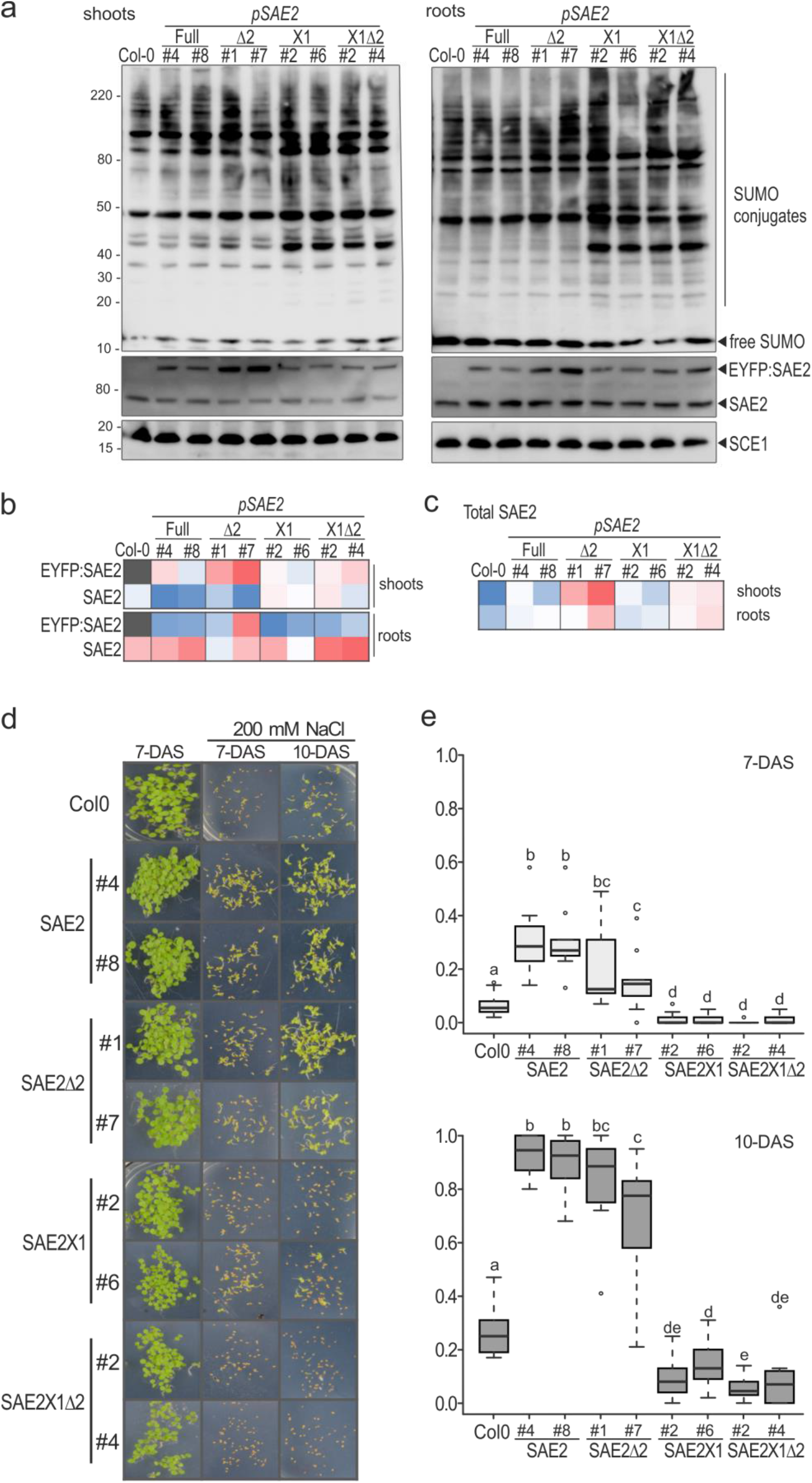
Effect of SAE2 cytosolic enrichment on SUMO conjugation and germination tolerance to salt stress. Two-independent transgenic plants expressing each of the following SAE2 proteoforms under the control of the pSAE2 promoter were generated: EYFP:SAE2 (#4 and #8), EYFP:SAE2Δ2 (#1 and #7), EYFP:SAE2X1 (#2 and #6), and EYFP:SAE2X1Δ2 (#2 and #4). These SAE2 proteoforms display different subcellular distribution as shown in Figure 2. (a) Effect of expression of SAE2 proteoforms on SUMO conjugates, SAE2 and SCE1 levels. Total protein extracts from shoots (left) and roots (right) from 11-day old seedlings were resolved by SDS-PAGE and examined by immunoblot analysis with anti-SUMO1 (top panel), anti-SAE2 (middle panel) and anti-SCE1 (bottom panel) antibodies. (b) Relative SAE2 proteoform levels (EYFP:SAE2 proteoforms and endogenous SAE2) were quantified from two-biological replicates and plotted in a heat map table were the highest level is in red, the lowest in blue and the median is in white. Shoots and roots values were treated as two independent groups. (c) Total SAE2 levels per sample were calculated by the addition of recombinant EYFP:SAE2 and endogenous SAE2 levels obtained in (b). (d) Seeds from plants the indicated transgenic plants were germinated on media in the absence or presence of 200 mM NaCl and germination rate scored at 7-days after sowing (DAS) or 10-DAS. Representative images are shown in (d). Germination was scored as cotyledon expansion and greening. Data was obtained from 10 biological replicates using 5 different seed batches (2 replicates form each seed batch). In each replicate, 40-50 seeds from each transgenic line were analyzed. Values obtained at 7-DAS (B) and 10-DAS (C) were plotted on the graphs. Data was analyzed by two-tailed T-test (unequal variance). Different letters denote statistically significant differences between groups (*P* < 0.05).

Similarly to SCE1 levels, we did not observe quantitative differences in global SUMO conjugates accumulation between transgenic and Col0 plants, suggesting that the achieved SAE2 levels did not reach a threshold to promote endogenous SUMO conjugation. In contrast, plants expressing SAE2 proteoforms enriched in the cytosol (the nucleo-cytosolic EYFP:SAE2Δ1L and EYFP:SAE2X1 forms; and the fully cytosolic EYFP:SAE2Δ1,2L and EYFP:SAE2X1Δ2 forms) displayed qualitative differences in the SUMO conjugate pattern (Figure 4a and Supplementary Figure 5). Although the identity of these substrates remain to be elucidated, we speculate that cytosolic SAE2 enrichment promotes SUMO conjugation of cytosolic substrates not accessible to the SUMO conjugation machinery in seedlings.

The effect of increased SAE2 levels on plant development varies depending on the SAE2 proteoform expressed. Rosette area size was only altered in plants expressing the partial cytoplasmic SAE2Δ1L proteoform under the control of the 35S promoter, which displayed significant smaller rosettes. On the contrary, SAE2 nuclear enrichment resulted in petiole elongation and this effect was reverted gradually by increasing accumulation of SAE2 in the cytosol, expressed under both promoters (Supplementary Figure 5). A similar effect was observed in flowering studies, while SAE2 nuclear enrichment resulted in a delay in flowering, this effect was reverted by the partial cytosolic enriched SAE2 forms (EYFP:SAE2Δ1L and EYFP:SAE2X1) (Supplementary Figure 6). No alterations in seed size or production were observed (Supplementary Figure 7), consistently with the initial observation showing that the endogenous SAE2, in seeds, is mainly present as the processed form (Figure 2e). This opposing effect between cytosolic-enriched and nuclear SAE2 proteoforms in plant growth was also observed in germination under salt stress (Figure 4 and Supplementary Figure 8). The tolerance conferred of nuclear SAE2 to salt stress is consistent with the protective role of the nuclear SUMO E3 ligase overexpression in homologous^28^ and heterologous systems^29,30^. Although technical developments are required to dissect the role of cytosolic SUMOylation, previous proteomic studies showed that cytosolic SUMO conjugates are quantitatively important^18^.

### Evolutionary acquisition by seed plants of the variable linker regions between NLS1 and NLS2 in the SAE2 C-terminal tail

For assessing the relevance in the plant kingdom of this uncovered regulatory mechanism, we assembled a curated dataset of *bona fide* SAE2 orthologous sequences from 42 representative land plant species by scanning different genome databases (Supplementary Table 1). Careful inspection of the sequence alignment at the SAE2 C-terminal tail reveals the occurrence of a linker region between NLS1 and NLS2 in seed plants, ranging in length from 32 amino acids in the model eudicot organism *A. thaliana* to 76 in the early seed gymnosperm plant *Gnetum montanum* (Supplementary Figure 9). Interestingly, while the linker region was also present in the gymnosperm living fossil *Ginkgo biloba*, the oldest extant representative of seed plants, it was absent from any non-seed plant (Figure 5 and Supplementary Figure 9), including the heterosporous ferns *Azolla filiculoides* and *Salvinia cucullata*, despite the fact that heterospory represents a key transition in the evolution of seed plants. This additional seed-specific SAE2 C-terminal linker region was poorly conserved at the length and amino acid composition level, except for a short stretch of around 10 acidic amino acid (mostly composed of aspartic acid residues) located near the NLS2 and interrupted by two up to five branched amino acids (Figure 5). Although a different origin of the last exon encoding for the SAE2 C-terminal tail region might account for the occurrence of this seed-specific C-tail linker region, examination of the last intron revealed its conservation both in terms of location and phase between seed and non-seed plants, such as mosses or lycophytes, (Supplementary Table 1 and Supplementary Figure 9). Altogether, our results support the independent acquisition of the linker region concomitantly with the emergence of seed plants. Although the exact precise role of this, mostly repetitive, linker region, it is at present unknown, it appears that repeated sequence patterns may provide regular spacing allowing optimal folding between protein functional domains and commonly overlap with intrinsically unstructured, low complexity or disordered regions (IURs) suggested to participate in protein-ligand binding, including protein-protein interactions^31–33^.

**Figure 5.**
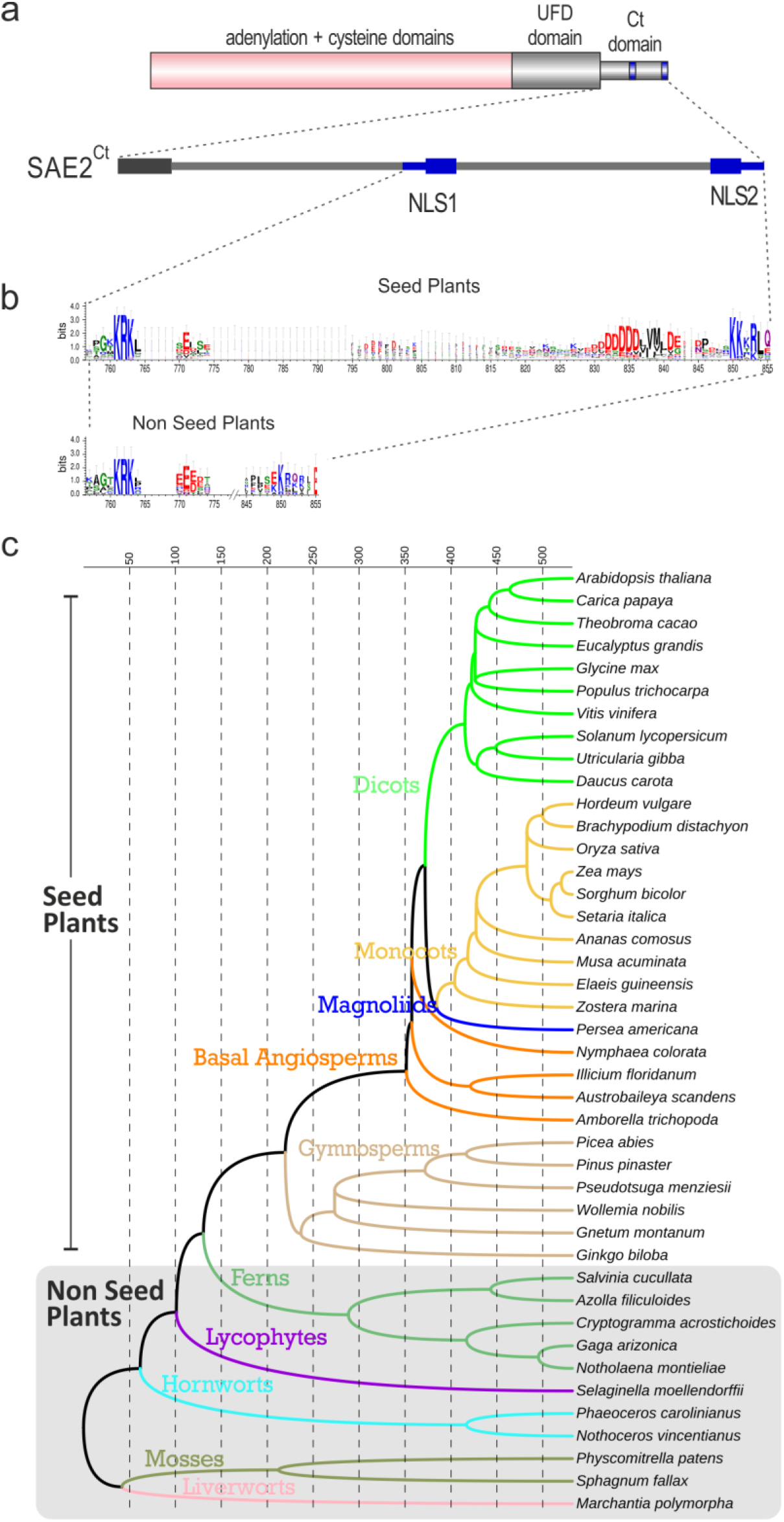
Conservation analysis of SAE2^Ct^ domain in plants. (a) Graphical representation of *Arabidopsis* SAE2 domains. (b) Logo representation of the multiple sequence alignments of the C-terminal tail region of 31 seed- and 11 non seed-plant SAE2 sequences. See Supplementary Fig. S9 for the original alignments. The overall height of the stack indicates the sequence conservation at that position, while the symbol height within the stack indicates the relative frequency of each amino acid within that position. (c) Taxonomic tree representing the evolutionary relationships among the 42 seed and non-seed land plant species. Topologies and divergence times were taken from TimeTree.

## Discussion

It is reported that SUMOylation machinery components display distinct subcellular distributions^6^ that may add precision to SUMO conjugation/deconjugation. Here, we propose that the subcellular distribution of the *Arabidopsis* SUMO activating enzyme E1 is dynamic. Although the predominant form is nuclear localized by means of two independent NLSs located at the E1 large subunit SAE2 C-terminal tail, limited proteolysis of the C-terminal NLS2 facilitates SAE2 accumulation in the cytosol. One of the remaining questions to be answered is at which subcellular compartment SAE2 processing takes place. A similar regulatory mechanism has been observed during adenovirus maturation, where proteolysis of the adenoviral protein VI C-terminus releases a NLS that results in a cytosolic enrichment and its function changes from a shuttling transport adapter to a viral structural protein^34^. Although limited proteolysis aiming to facilitate surface exposure of a cryptic NLS, or to generate a soluble NLS-containing fragment, is a well-known mechanism to promote nuclear import of a cytosolic protein^35^, our results, together with others, suggest that proteolytic cleavage of NLS constitutes an emerging mechanism to facilitate nuclear exclusion.

Considering that no <u>n</u>uclear <u>e</u>xported <u>s</u>ignals (NES) has been identified in SAE2, we favor the hypothesis that SAE2 processing may occur at the nucleus, resulting in a reduction of its nuclear retention. In mammals, SUMO E1 subcellular localization is also regulated at different levels. The NLS present in the human E1 large subunit Uba2 is the only signal required for the E1 efficient import into the nucleus^25^ and, an additional post-translational modification by SUMO at the Uba2 C-terminal domain contributes to its nuclear retention^36^. Overall, regulation of E1 subcellular distribution seems to be conserved across evolutionary kingdoms, although through distinct molecular mechanisms.

We propose that SAE2 cytosolic enrichment would constitute a molecular switch to promote cytosolic SUMOylation, which, according to our results, counteracts the effect of nuclear SUMOylation on plant growth and salt stress tolerance during germination. These opposing roles have also been proposed to occur in cardiomyocites, where nuclear SUMOylation is detrimental to heart function while extra-nuclear SUMOylation is cardioprotective, suggesting that compartmentalization of SUMO conjugation could account for the capability to rapidly switch between cell physiological states^37^.

We have shown that SAE2 processing is tissue specific, being predominant in seeds. It is well described the important role of SUMO during seed development. SUMO1/2, SAE2 and SCE1 T-DNA insertion mutants are lethal at the embryonic stage; and the mutant of the E3 ligase SIZ1 and HYP1-2 shows a 33,2% and 6,9% aborted seeds, respectively. Our findings constitute an additional layer of complexity by pointing to an important role of cytosolic SUMOylation in the transition from the embryo growth phase to seed maturation phase. Consistent with this potential specialized role, the *Arabidopsis* SAE2 C-terminal tail structural signature / linker region is only found in seed plants. Further research aiming to identify the protease responsible for SAE2 proteolytic processing will be key to deeply understand the biological consequences of SAE2 cytosolic enrichment and its role in seed appearance in evolution.

## Methods

### Plant Material and Growth Conditions

For *in vitro* cultures, seeds were stratified for 3 days, plated on Murashige and Skoog salts (Duchefa), pH 5.7, supplemented with 0.8% BactoAgar (Difco), and transferred to a tissue culture room in a LD photoperiod (16 h light/8 h dark) at 22 °C. For soil cultures, plants were grown in growth chambers in a LD photoperiod at 22°C. For salt stress experiments, *in vitro* media was supplemented with 200 mM NaCl.

### Protein extraction and immunoblot

Plant tissue was ground in liquid nitrogen and proteins extracted with 50 mM Tris-HCl pH 8, 150 mM NaCl, 0.2% Triton X-100, 1 mM PMSF, 1 μg ml^−1^ pepstatin, 1 μg ml^−1^ leupeptin, 2 mM N-ethylmaleimide, 10 mM iodoacetamide and 5 mM EDTA (extraction buffer). In seeds and siliques, total proteins were extracted with urea buffer (100 mM Tris-HCl pH 7.5, 8 M urea, 0.20% Triton X-100, 0.20% Sakosyl, 1 mM PMSF, 1 μg ml^−1^ pepstatin, 1 μg ml^−1^ leupeptin, 2 mM N-ethylmaleimide, 10 mM iodoacetamide and 5 mM EDTA). 18 μg of total protein were resolved under reducing conditions by using SDS polyacrylamide gels and NuPage Novex 4–12% Bis/Tris Gels (Invitrogen) for SUMO conjugates analysis or 10% for SAE2 immunodetection. Proteins were transferred onto polyvinylidene difluoride (PVDF) membranes (Millipore), incubated with anti-SUMO1^14^ or anti-SAE2^15^ antibodies overnight followed by incubation with secondary antibody, peroxidase-conjugated anti-rabbit (GE Healthcare), for 1 h at room temperature in TBST (20 mM Tris-HCl, pH 7.6, 20 mM NaCl, 0.1% (v/v) Tween20) supplemented with 3% non-fat dry milk. Peroxidase activity was developed in ECL Plus reagent (GE Healthcare) and chemiluminescence signal captured with the LAS-4000 imaging system (Fujifilm).

### SAE2 proteolytic processing analysis

SAE2^UFDCt^ was cloned into cloned into pET28a (Novagene) to encode a N-terminal Histidine tag and produced as previously described^38^. 350μg of freshly protein extracts from rosette and silique Col-0 plants were incubated with 10μg of recombinant His:SAE2^UFDCt^ protein in the presence of the extraction buffer supplemented with ATP. The control reaction was done by the incubating of the His:UFDCt without any plant extract. Reaction mixtures were incubated during 60 min at 30°C. After the specified incubation times, reactions were stopped by the addition of protein loading buffer and boiled for 10 min. 15-fold dilution of the result sample was loaded and resolved in 12% SDS-PAGE followed by immunoblot analysis anti-polyHIS monoclonal antibodies (Sigma, H1020). Scale-up experiments were done in order to purify His:SAE2^UFDCt^ and perform subsequent MS analysis. Scale-up conditions: 100μg of His:SAE2^UFDCt^ incubated with 3.5mg of rosette or siliques protein extracts from Col-0 plants during 60 min at 30°C. Reaction products were Ni^2+^-affinity purified and subjected to mass spectrometry analysis.

### Transient Expression of Fluorescent Protein Fusions in Onion Cells

*SAE2 and SAE2*^*UFDCt*^ proteoforms were fused in frame to the 3’ end of the coding sequences of yellow fluorescent protein (YFP) or cyan fluorescent protein (CFP), respectively, downstream of the 35S constitutive promoter. Onion epidermal cells were bombarded with 5 μg of each DNA construct using a helium biolistic gun (BIO-RAD). Treated epidermal cells were kept in the dark at room temperature for 16 h before analysis by confocal microscopy (Confocal Olympus FV 1000). YFP was excited with a 515-nm argon laser and images collected with a 550- to 630-nm range. Image analysis and data quantification was performed as previously described^39^.

### Importin binding assays

EYFP:SAE2^Ct^, EYFP:NLS1 and EYFP:NLS2 were fused in frame to the 3’ end of a 6xHis:2x-flag tag into the pET28a bacteria expression vector. Recombinant proteins were produced as previously described^38^. 1 μM of the EYFP fusion variants were incubated in the presence of increasing concentrations (0, 0.5, 1.5 or 4 μM) of native α-importin (M0) or harboring mutations in the major groove (M1) or the minor groove (M2) for NLS binding. EYFP was used as a negative control. Protein mixtures were incubated in 20 mM HEPES-NaOH pH 7.4, 0.1 M NaCl, 2 mM DTT, 12.5% glycerol and 0.1% BSA at room temperature for 1h. Protein complexes were resolved by native 7.5% PAGE in Tris-Glycine running buffer containing 1 mM DTT and 10% glycerol for 90 min at 160V and 4°C. Fluorescence signal from resolved products was captured with LAS-4000 imaging system (Fujifilm) using a green filter.

### Bioinformatics analyses

We generated a curated dataset of SAE2 protein sequences from 42 representative land plant species by scanning their proteomes with BLAST using the *A. thaliana* SAE2 protein sequence as query (Supplementary Table 1). Only full-length or nearly full-length sequences matching canonical SAE2 INTERPRO functional domains were considered. For species displaying lineage specific duplicated *SAE2* genes, the copy resulting in the best BLAST reciprocal hit was considered as true co-ortholog in further analysis. Multiple amino acid sequence alignments were obtained using Muscle^40^. Graphical representation of multiple sequence alignments (logos) were reconstructed using WebLogo 3.7.4 software (http://weblogo.threeplusone.com/^41^). A taxonomic tree representing the evolutionary relationships among the 42 selected plant species was constructed using topologies and divergence times from TimeTree^42^.

## Supporting information

sequence codes

## Acknowledgements

We thank the technical support from members of the Greenhouse and Microscopy facilities at CRAG. We greatly thank Cristina Cañadas for technical support at LML lab. Importin constructs were kindly provided by Dr. Shunichi Kosugi. This work was supported by European Research Council (ERC-2007-StG-205927) and FEDER/Ministerio de Ciencia, Innovación y Universidadese (BIO2017-89874-R) grants to LML. A.M. was supported by predoctoral fellowships FPU12/05292, and L.C.M. by research contracts through the CRAG. We acknowledge financial support from the Generalitat de Catalunya (Xarxa de Referència en Biotecnologia and 2017SGR1211) and the Spanish Ministry of Economy and Competitiveness, through the “Severo Ochoa Programme for Centres of Excellence in R&D” 2016-2019 (SEV-2015-0533).

## References

1. Deribe, Y. L., Pawson, T. & Dikic, I. Post-translational modifications in signal integration. Nat. Struct. Mol. Biol. 17, 666–672 (2010).

2. Rogers, L. D. & Overall, C. M. Proteolytic post-translational modification of proteins: proteomic tools and methodology. Mol. Cell. Proteomics 12, 3532–3542 (2013).

3. Nacerddine, K. et al. The SUMO pathway is essential for nuclear integrity and chromosome segregation in mice. Dev. Cell 9, 769–779 (2005).

4. Saracco, S. A., Miller, M. J., Kurepa, J. & Vierstra, R. D. Genetic analysis of SUMOylation in Arabidopsis: conjugation of SUMO1 and SUMO2 to nuclear proteins is essential. Plant Physiol. 145, 119–134 (2007).

5. Bentsink, L. & Koornneef, M. Seed Dormancy and Germination. Arab. B. e0119 (2008) doi:10.1199/tab.0119.

6. Benlloch, R. & Lois, L. M. Sumoylation in plants: Mechanistic insights and its role in drought stress. J. Exp. Bot. 69, 4539–4554 (2018).

7. Augustine, R. C. & Vierstra, R. D. SUMOylation: re-wiring the plant nucleus during stress and development. Curr. Opin. Plant Biol. 45, 143–154 (2018).

8. Verma, V., Croley, F. & Sadanandom, A. Fifty shades of SUMO: its role in immunity and at the fulcrum of the growth-defence balance. Mol. Plant Pathol. 19, 1537–1544 (2018).

9. Castro, P. H., Tavares, R. M., Bejarano, E. R. & Azevedo, H. SUMO, a heavyweight player in plant abiotic stress responses. Cell. Mol. Life Sci. 69, 3269–3283 (2012).

10. Rosa, M. T. & Abreu, I. A. Exploring the regulatory levels of SUMOylation to increase crop productivity. Curr Opin Plant Biol 49, 43–51 (2019).

11. Sriramachandran, A. M. & Dohmen, R. J. SUMO-targeted ubiquitin ligases. Biochim. Biophys. Acta - Mol. Cell Res. 1843, 75–85 (2014).

12. Lois, L. M. & Lima, C. D. Structures of the SUMO E1 provide mechanistic insights into SUMO activation and E2 recruitment to E1. EMBO J. 24, 439–451 (2005).

13. Hickey, C. M., Wilson, N. R. & Hochstrasser, M. Function and regulation of SUMO proteases. Nat. Rev. Mol. Cell Biol. 13, 755–766 (2012).

14. Castaño-Miquel, L., Seguí, J. & Lois, L. M. M. Distinctive properties of Arabidopsis SUMO paralogues support the in vivo predominant role of AtSUMO1/2 isoforms. Biochem. J. 436, 581–590 (2011).

15. Castaño-Miquel, L. et al. Diversification of SUMO-activating enzyme in arabidopsis: Implications in SUMO conjugation. Mol. Plant 6, 1646–1660 (2013).

16. Morrell, R. & Sadanandom, A. Dealing With Stress: A Review of Plant SUMO Proteases. Front. Plant Sci. 10, (2019).

17. Hammoudi, V., Vlachakis, G., Schranz, M. E. & van den Burg, H. A. Whole-genome duplications followed by tandem duplications drive diversification of the protein modifier SUMO in Angiosperms. New Phytol. 211, 172–185 (2016).

18. Rytz, T. C. et al. SUMOylome profiling reveals a diverse array of nuclear targets modified by the SUMO ligase SIZ1 during heat stress. Plant Cell 30, 1077–1099 (2018).

19. Tomanov, K. et al. Arabidopsis PIAL1 and 2 promote SUMO chain formation as E4-type SUMO ligases and are involved in stress responses and sulfur metabolism. Plant Cell 26, 4547–4560 (2014).

20. Reiland, S. et al. Large-scale Arabidopsis phosphoproteome profiling reveals novel chloroplast kinase substrates and phosphorylation networks. Plant Physiol. 150, 889–903 (2009).

21. Meyer, L. J., Gao, J., Xu, D. & Thelen, J. J. Phosphoproteomic analysis of seed maturation in Arabidopsis, rapeseed, and soybean. Plant Physiol. 159, 517–528 (2012).

22. Nakagami, H. et al. Large-scale comparative phosphoproteomics identifies conserved phosphorylation sites in plants. Plant Physiol. 153, 1161–1174 (2010).

23. Nukarinen, E., Tomanov, K., Ziba, I., Weckwerth, W. & Bachmair, A. Protein sumoylation and phosphorylation intersect in Arabidopsis signaling. Plant J. 91, 505–517 (2017).

24. Castaño-Miquel, L. et al. SUMOylation Inhibition Mediated by Disruption of SUMO E1-E2 Interactions Confers Plant Susceptibility to Necrotrophic Fungal Pathogens. Mol. Plant 10, 709–720 (2017).

25. Moutty, M. C., Sakin, V. & Melchior, F. Importin alpha/beta mediates nuclear import of individual SUMO E1 subunits and of the holo-enzyme. Mol. Biol. Cell 22, 652–660 (2011).

26. Kosugi, S. et al. Six classes of nuclear localization signals specific to different binding grooves of importin alpha. J Biol Chem 284, 478–485 (2009).

27. Ito, J. et al. Analysis of the Arabidopsis cytosolic proteome highlights subcellular partitioning of central plant metabolism. J. Proteome Res. 10, 1571–1582 (2011).

28. Miura, K. & Nozawa, R. Overexpression of SIZ1 enhances tolerance to cold and salt stresses and attenuates response to abscisic acid in Arabidopsis thaliana. Plant Biotechnol. 31, 167–172 (2014).

29. Mishra, N. et al. Overexpression of the Rice SUMO E3 Ligase Gene OsSIZ1 in Cotton Enhances Drought and Heat Tolerance, and Substantially Improves Fiber Yields in the Field under Reduced Irrigation and Rainfed Conditions. Plant Cell Physiol. 58, 735–746 (2017).

30. Mishra, N., Srivastava, A. P., Esmaeili, N., Hu, W. & Shen, G. Overexpression of the rice gene OsSIZ1 in Arabidopsis improves drought-, heat-, and salt-tolerance simultaneously. PLoS One 13, e0201716 (2018).

31. Zhou, Y., Liu, J., Han, L., Li, Z.-G. & Zhang, Z. Comprehensive analysis of tandem amino acid repeats from ten angiosperm genomes. BMC Genomics 12, 632 (2011).

32. Kumar, A. S., Sowpati, D. T. & Mishra, R. K. Single Amino Acid Repeats in the Proteome World: Structural, Functional, and Evolutionary Insights. PLoS One 11, e0166854 (2016).

33. Simon, M. & Hancock, J. M. Tandem and cryptic amino acid repeats accumulate in disordered regions of proteins. Genome Biol. 10, R59 (2009).

34. Wodrich, H. et al. Switch from capsid protein import to adenovirus assembly by cleavage of nuclear transport signals. EMBO J. 22, 6245–6255 (2003).

35. Boulikas, T. Nuclear localization signals (NLS). Crit. Rev. Eukaryot. Gene Expr. 3, 193–227 (1993).

36. Truong, K., Lee, T. D., Li, B. & Chen, Y. Sumoylation of SAE2 C terminus regulates SAE nuclear localization. J. Biol. Chem. 287, 42611–42619 (2012).

37. Le, N.-T., Martin, J. F., Fujiwara, K. & Abe, J. Sub-cellular localization specific SUMOylation in the heart. Biochim. Biophys. Acta - Mol. Basis Dis. 1863, 2041–2055 (2017).

38. Castaño-Miquel, L. & Lois, L. M. Kinetic Analysis of Plant SUMO Conjugation Machinery. in Methods in Molecular Biology vol. 1450 107–123 (Humana Press Inc., 2016).

39. Mas, A., Amenós, M. & Lois, L. M. Quantitative Analysis of Subcellular Distribution of the SUMO Conjugation System by Confocal Microscopy Imaging. in Methods in Molecular Biology vol. 1450 135–150 (Humana Press Inc., 2016).

40. Edgar, R. C. MUSCLE: multiple sequence alignment with high accuracy and high throughput. Nucleic Acids Res 32, 1792–1797 (2004).

41. Crooks, G. E., Hon, G., Chandonia, J. M. & Brenner, S. E. WebLogo: A sequence logo generator. Genome Res. 14, 1188–1190 (2004).

42. Kumar, S., Stecher, G., Suleski, M. & Hedges, S. B. TimeTree: A Resource for Timelines, Timetrees, and Divergence Times. Mol. Biol. Evol. 34, 1812–1819 (2017).

