## supplementary figures for "Evolution of molecular determinants for SUMO-activating enzyme subcellular localization in plants"

### Supplementary information

**Supplementary Figure 1.**

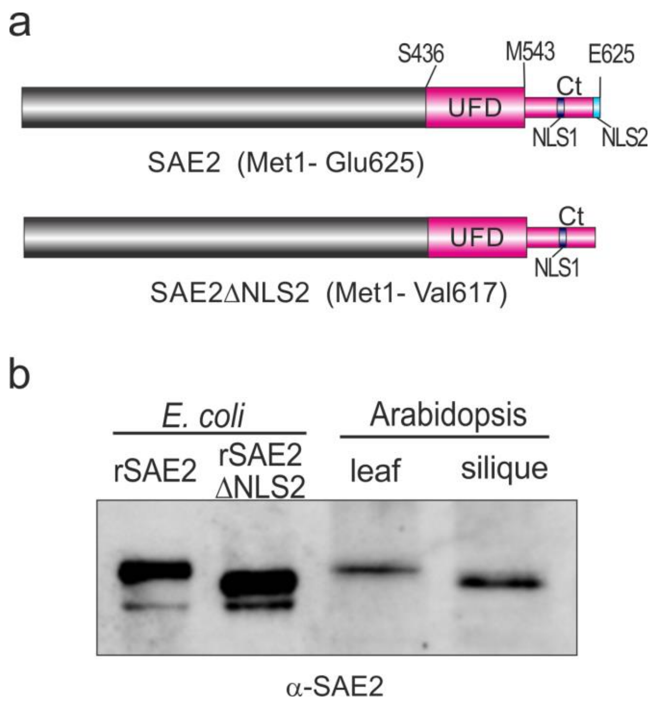


**Supplementary Figure 1. Production of recombinant SAE2 proteoforms as standards for SDS-PAGE SAE analysis**.

(a) Full length SAE2 (M1-E625) and its deletion variant SAE2ΔNLS2 (M1-V617) were cloned into pET28a expression vector at 5’ Nco I site and 3’ NotI or SacI. As a result, the translation products did not contain any additional amino acid from the plasmid.

(b) Total protein extracts from 5-week old Col-0 rosette leaves and mature green siliques, and *E. coli* protein extracts containing the generated recombinant SAE2 standards were analyzed by immunoblotting with anti-SAE2 antibodies. The comparative analysis indicated that SAE2 present in rosette leaves displays the same apparent MW as the recombinant full length SAE2, while SAE2 present in siliques displays an apparent MW consistent with the SAE2 deletion variant SAE2ΔNLS2 (M1-V617).

**Supplementary Table 1. DNA constructs generated.**

| Construct | Code | Cloning vector | Insert | Fusion tag | Plant selection marker | Bacteria selection marker |
|---|---|---|---|---|---|---|
| pSAE2::EYFP:SAE2 *XNLS1* | pLC01 | pH7m24gw.3 | SAE2 (Met1-Glu625) K582T K583G | EYFP | Hygromycin | |
| pSAE2::EYFP:SAE2 *XNLS1ΔNLS2* | pLC02 | pH7m24gw.3 | SAE2 (Met1-Val617) K582T K583G | EYFP | Hygromycin | |
| pSAE2::EYFP:SAE2 Full | pLC23 | pH7m24gw.3 | SAE2 (Met1-Glu625) | EYFP | Hygromycin | |
| pSAE2::GFP-GUS | pLC03 | pH7m24gw.3 | -1495..-1 genomic | GFP-GUS | Hygromycin | |
| pB2GW7::EYFP:SAE2Full | pLC29 | pB2GW7 | SAE2 (Met1-Glu625) | EYFP | PPT | spectinomycin |
| pB2GW7::EYFP:SAE2 *ΔNLS2* | pLC30 | pB2GW7 | SAE2 (Met1-Val617) | EYFP | PPT | spectinomycin |
| pB2GW7::EYFP:SAE2 *ΔNLS1,linker* | pLC31 | pB2GW7 | SAE2 (Met1-Ser574)+(Ser618-Glu625) | EYFP | PPT | spectinomycin |
| pB2GW7::EYFP:SAE2 *ΔNLS1,2linker* | pLC32 | pB2GW7 | SAE2 (Met1-Ser574) | EYFP | PPT | spectinomycin |
| pWEN25::SAE2 *ΔNLS1,linker* | pLC11 | pWEN25 | SAE2 (Met1-Ser574)+(Ser618-Glu625) | EYFP | | Ampicillin |
| pWEN25::SAE2 *ΔNLS1,2linker* | pLC12 | pWEN25 | SAE2 (Met1-Ser574) | EYFP | | Ampicillin |
| pWEN25::SAE2 *ΔNLS2* | pLC13 | pWEN25 | SAE2 (Met1-Val617) | EYFP | | Ampicillin |
| pWEN25::SAE2 Full | pLC14 | pWEN25 | SAE2 (Met1-Glu625) | EYFP | | Ampicillin |
| pWEN25::SAE2 *XNLS1* | pLC24 | pWEN25 | SAE2 (Met1-Glu625) K582T K583G | EYFP | | Ampicillin |
| pWEN25::SAE2 *Δlinker* | pLC25 | pWEN25 | SAE2 (Met1-Leu586)+(Ser618-Glu625) | EYFP | | Ampicillin |
| pWEN25::SAE2 *XNLS1ΔNLS2* | pLC35 | pWEN25 | SAE2 (Met1-Val617) K582T K583G | EYFP | | Ampicillin |
| pWEN25::UFDCt | pJS37 | pWEN25 | SAE2 (Ser436-Glu625) | EYFP | | Ampicillin |
| pWEN25::UFD | pJS36 | pWEN25 | SAE2 (Ser436-Met543) | EYFP | | Ampicillin |
| pWEN25::EYFP:Ct | pCCM10 | pWEN25 | SAE2 (Met543-Glu625) | EYFP | | Ampicillin |
| pWEN25::EYFP:NLS1 | pCCM13 | pWEN25 | SAE2 (Pro579-Leu586) | EYFP | | Ampicillin |
| pWEN25::EYFP:NLS2 | pCCM14 | pWEN25 | SAE2 (Ser618-Glu625) | EYFP | | Ampicillin |
| pWEN25::UFDCt *ΔNLS2* | pLC15 | pWEN25 | SAE2 (Ser436-Val617) | EYFP | | Ampicillin |
| pET28b::AtUFDCt | pJS033 | pET28b | SAE2 (Ser436-Glu625) | His | | Kanamycin |
| pET28a::flag:EYFP:Ct | pAMG01 | pET28a | SAE2 (Met543-Glu625) | His:flag:EYFP | | Kanamycin |
| pET28a::flag:EYFP:NLS1 | pAMG02 | pET28a | SAE2 (Pro579-Leu586) | His:flag:EYFP | | Kanamycin |
| pET28a::flag:EYFP:NLS2 | pAMG03 | pET28a | SAE2 (Ser618-Glu625) | His:flag:EYFP | | Kanamycin |
| pET28a::flag:EYFP | pAMG04 | pET28a | | His:flag:EYFP | | Kanamycin |
| pET28a::SAE2 | | pET28a | SAE2 (Met1-Glu625) | none | | Kanamycin |
| pET28a::SAE2ΔNLS2 | pCCM36 | pET28a | SAE2 (Met1-Val617) | none | | Kanamycin |

**Supplementary Figure 2.**

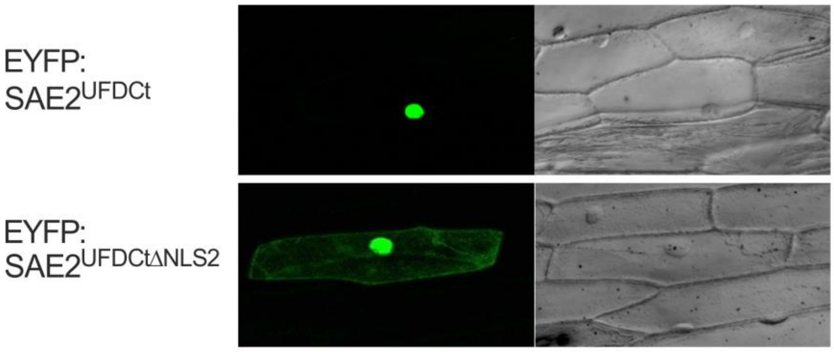


**Supplementary Figure 2. NLS2 is essential for complete nuclear retention of SAE2$^{UFDCt}$.** SAE2$^{UFDCt}$ domain (Ser436-Glu625) or the $\Delta$NLS2 SAE2$^{UFDCt}$ variant (Ser436-Val617) were fused to EYFP and transiently expressed in onion epidermal cells. Light transmission images of cells expressing the recombinant proteins are shown next to the corresponding fluorescence image.

**Supplementary Figure 3.**

a

| Name | Vector | Promoter | Tag | SAE2 variant | functional NLS | resistance | transgenic line codes |
|---|---|---|---|---|---|---|---|
| **Full** | pB2GW7 | p35S | EYFP | 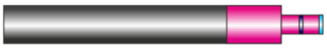 | NLS1<br>NLS2 | PPT | Full.8<br>Full.10 |
| **Δ2** | pB2GW7 | p35S | EYFP | 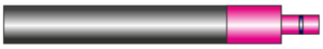 | NLS1 | PPT | Δ2.5<br>Δ2.8 |
| **Δ1L** | pB2GW7 | p35S | EYFP | 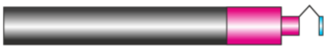 | NLS2 | PPT | Δ1L.4<br>Δ1L.5 |
| **Δ1,2L** | pB2GW7 | p35S | EYFP | 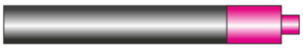 | - | PPT | Δ1,2L.8<br>Δ1,2L.10 |

b

| Name | Vector | Promoter | Tag | SAE2 variant | functional NLS | resistance | transgenic line codes |
|---|---|---|---|---|---|---|---|
| **Full** | pH7m24g w.3 | pSAE2 | EYFP | 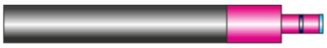 | NLS1<br>NLS2 | Hyg | Full.4<br>Full.8 |
| **Δ2** | pH7m24g w.3 | pSAE2 | EYFP | 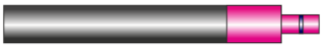 | NLS1 | Hyg | Δ2.1<br>Δ2.7 |
| **X1** | pH7m24g w.3 | pSAE2 | EYFP | 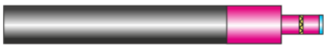 | NLS2 | Hyg | X1.2<br>X1.6 |
| **X1Δ2** | pH7m24g w.3 | pSAE2 | EYFP | 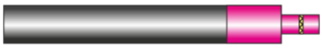 | - | Hyg | X1Δ2.2<br>X1Δ2.4 |

**Supplementary Figure 3.** Description of the transgenic lines generated. Expression of the SAE2 variant was driven under the control of the p35S promoter (a) or the pSAE2 promoter (b).

**Supplementary Figure 4.**

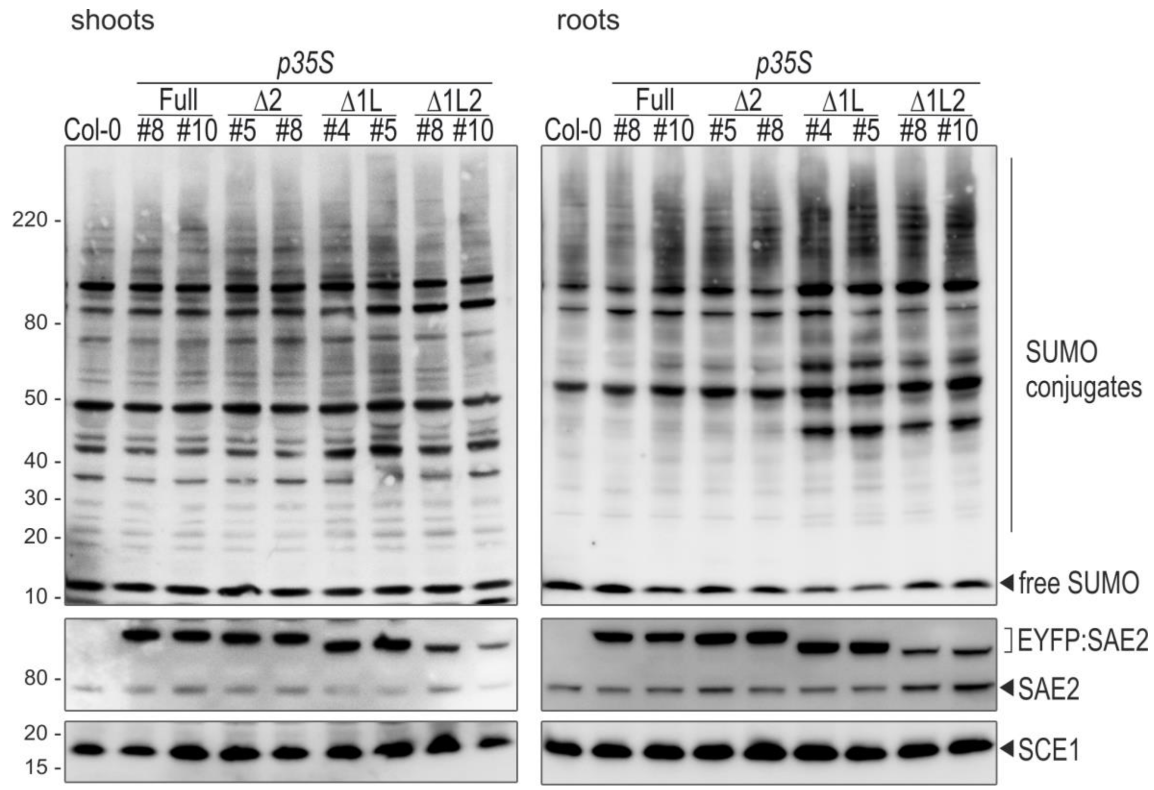


**Supplementary Figure 4. Molecular characterization of SUMO conjugation system in plants expressing SAE2 proteoforms under the control of the p35S promoter.**

Two-independent transgenic plants expressing each of the following SAE2 proteoforms under the control of the p35S promoter were generated: EYFP:SAE2 (#8 and #10), EYFP:SAE2Δ2 (#5 and #8), EYFP:SAE2Δ1L (#4 and #5), and EYFP:SAE2Δ1,2L (#8 and #10). These SAE2 proteoforms display different subcellular distribution as shown in Figure 2.

The effect of expression of SAE2 proteoforms on SUMO conjugates, SAE2 and SCE1 levels is shown. Total protein extracts from shoots (left) and roots (right) from 11-day old seedlings were resolved by SDS-PAGE and examined by immunoblot analysis with anti-SUMO1 (top panel), anti-SAE2 (middle panel) and anti-SCE1 (bottom panel) antibodies.

**Supplementary Figure 5.**

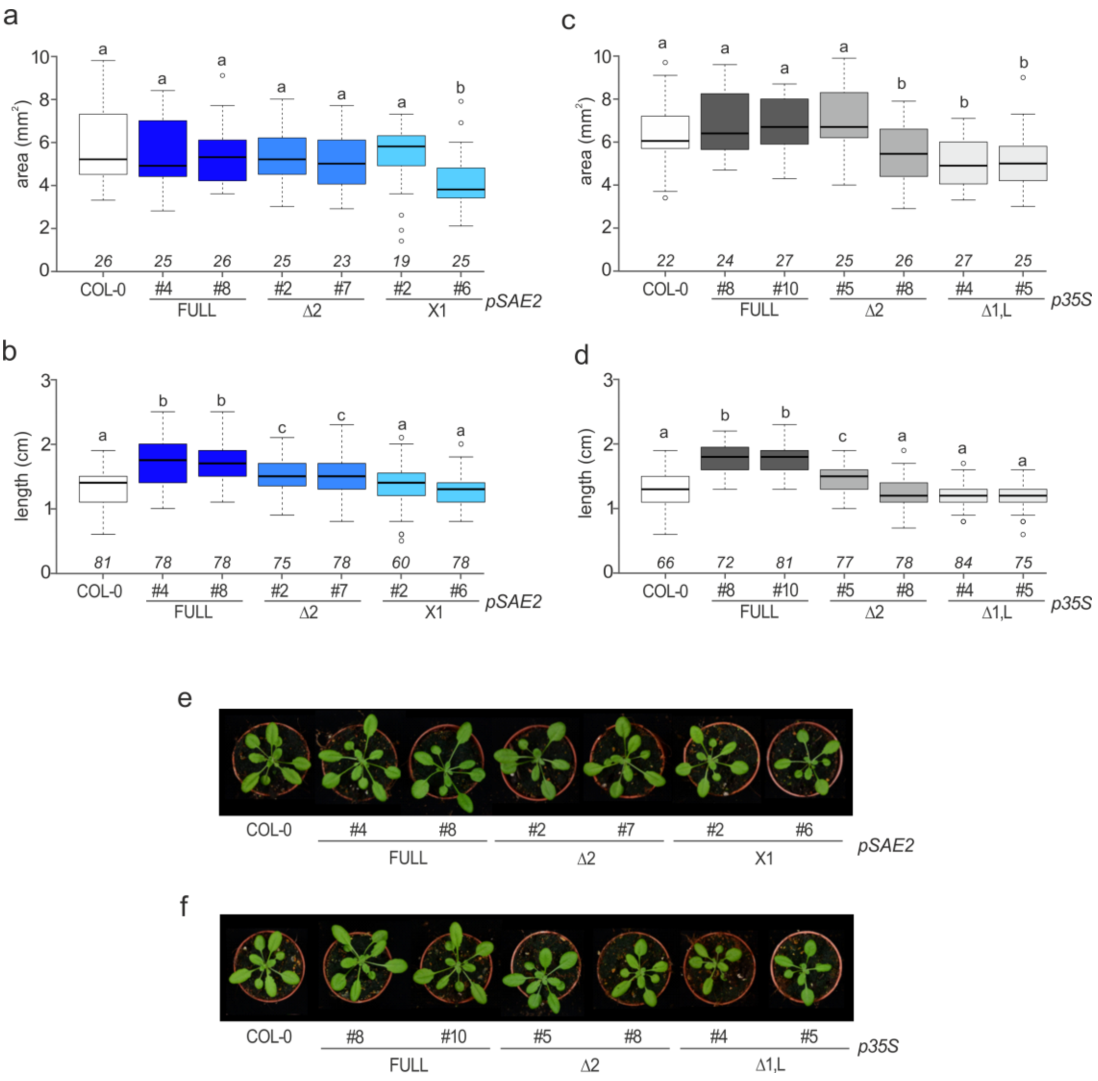


**Supplementary Figure 5.** Effect of SAE2 proteoforms expression on rosette size.

For the analysis, we used 21-old plants expressing the indicated SAE2 proteoforms under the control of the pSAE2 (a, b) or the p35S (c, d) promoters, grown under LD photoperiod and 22ºC and four biological replicates were performed. Images of representative plants are shown (e, f).

Measures were obtained by selecting the rosette area with the freehand tool included in the ImageJ software (a, c). Petiole length was measured on the rosette leaves at positions 3, 4 and 5 by using ImageJ (b, c). Data was plotted on the graphs and analyzed by two-tailed T-test. Different letters denote statistically significant differences between groups ($P < 0.05$).

**Supplementary Figure 6.**

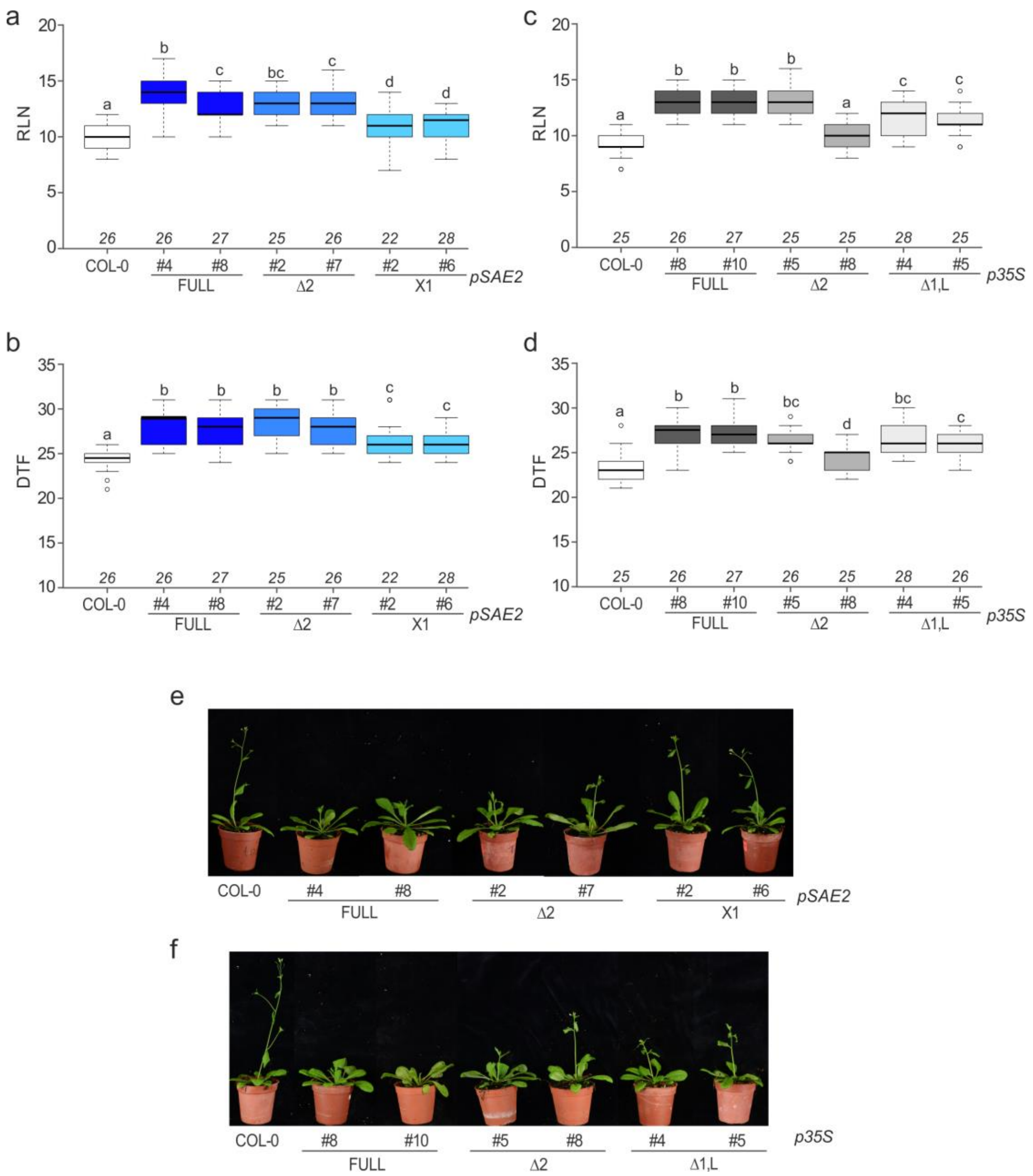


**Supplementary Figure 6.** Effect of SAE2 proteoforms expression on flowering.

For the analysis, we used 4-week old plants expressing the indicated SAE2 proteoforms under the control of the pSAE2 (a, b) or the p35S (c, d) promoters, grown under LD photoperiod and 22ºC and four biological replicates were performed. Images of representative plants are shown (e, f).

Rosette leaf number (RLN) at flowering and days to flowering (DTF) were scored when the inflorescence had reached 1 cm. Data was plotted on the graphs and analyzed by two-tailed T-test. Different letters denote statistically significant differences between groups ($P < 0.05$).

**Supplementary Figure 7.**

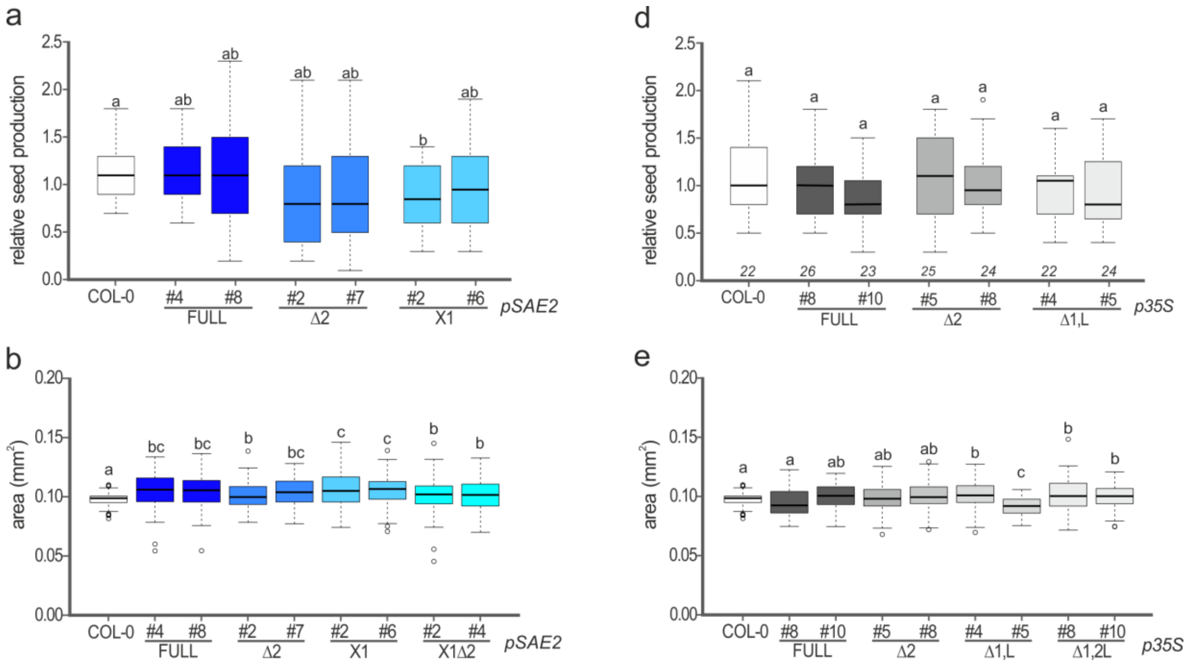


**Supplementary Figure 7.** Effect of SAE2 proteoforms expression on seed production and size.

For the analysis, we used four different seed batches from plants expressing the indicated SAE2 proteoforms under the control of the pSAE2 (a, b) or the p35S (c, d) promoters, grown under LD photoperiod and 22ºC. Relative seed production was calculated by weighting total seeds obtained per plant from 5-7 plants per biological replicate (a, d). Seed size was measured with SmartGrain software (b, e) (Takanari Tanabata, Taeko Shibaya, Kiyosumi Hori, Kaworu Ebana, and Masahiro Yano, "*SmartGrain*: High-throughput phenotyping software for measuring seed shape through image analysis", ***Plant Physiology***, Vol.160(4), pp.1871-1880. doi: http://dx.doi.org/10.1104/pp.112.205120). Data was plotted on the graphs and analyzed by two-tailed T-test. Different letters denote statistically significant differences between groups (*P* < 0.05).

**Supplementary Figure 8.**

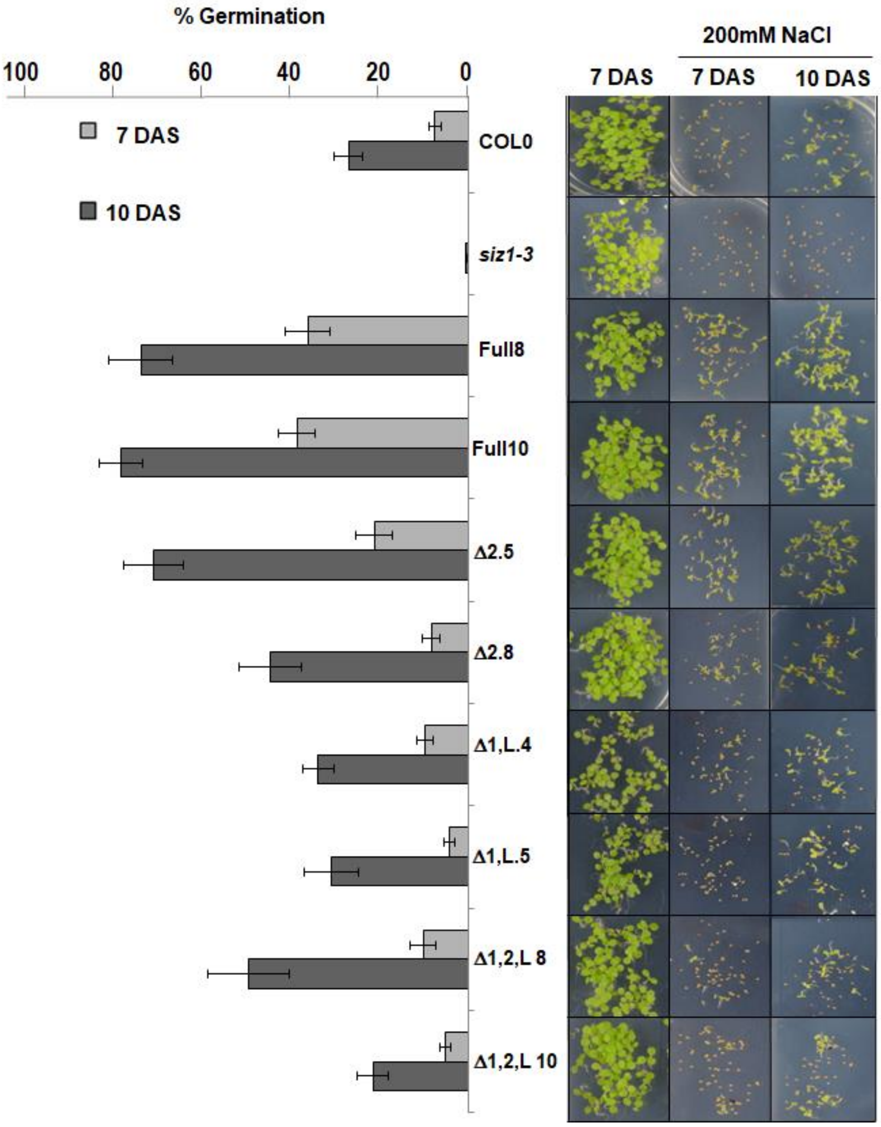


**Supplementary Figure 8.** Effect of SAE2 proteoforms constitutive expression in germination under salt stress.

Seeds from plants the indicated transgenic plants were germinated on media in the absence or presence of 200 mM NaCl and germination rate scored at 7-days after sowing (DAS) or 10-DAS. Representative images are shown in (d). Germination was scored as cotyledon expansion and greening. Data was obtained from 10 biological replicates using 5 different seed batches (2 replicates form each seed batch). In each replicate, 40-50 seeds from each transgenic line were analyzed. Values obtained at 7-DAS (B) and 10-DAS (C) were plotted on the graphs.

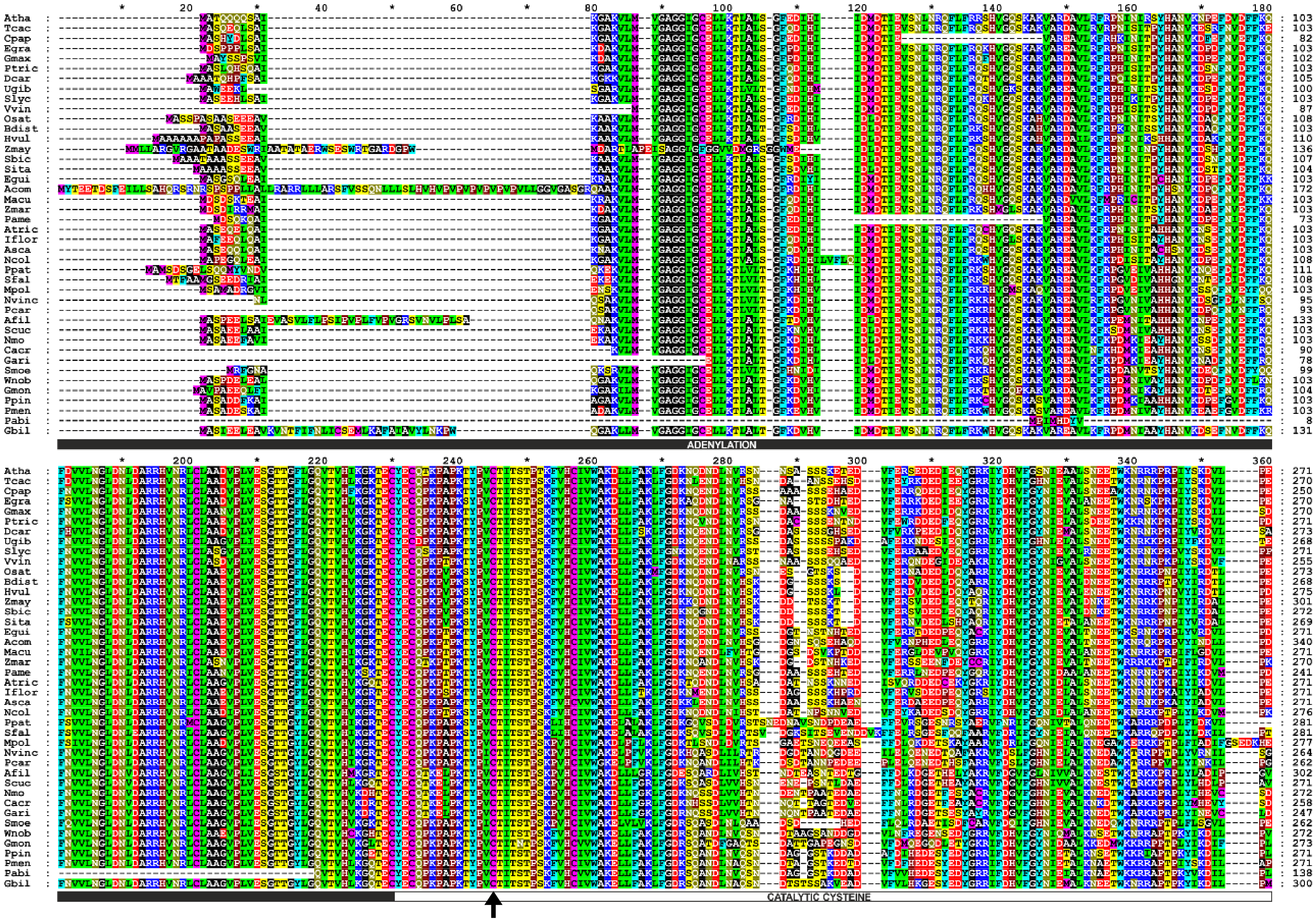
Atha
Tcac
Cpap
Egra
Gmax
Ptric
Dcar
Ugib
Slyc
Vvin
Osat
Bdist
Hvul
Zmay
Sbic
Sita
Egui
Acom
Macu
Zmar
Pame
Atric
Iflor
Asca
Ncol
Ppat
Sfal
Mpol
Nvinc
Pcar
Afil
Scuc
Nmo
Cacr
Gari
Smoe
Wnob
Gmon
Ppin
Pmen
Pabi
Gbil
ADENYLATION
CATALYTIC CYSTEINE

03

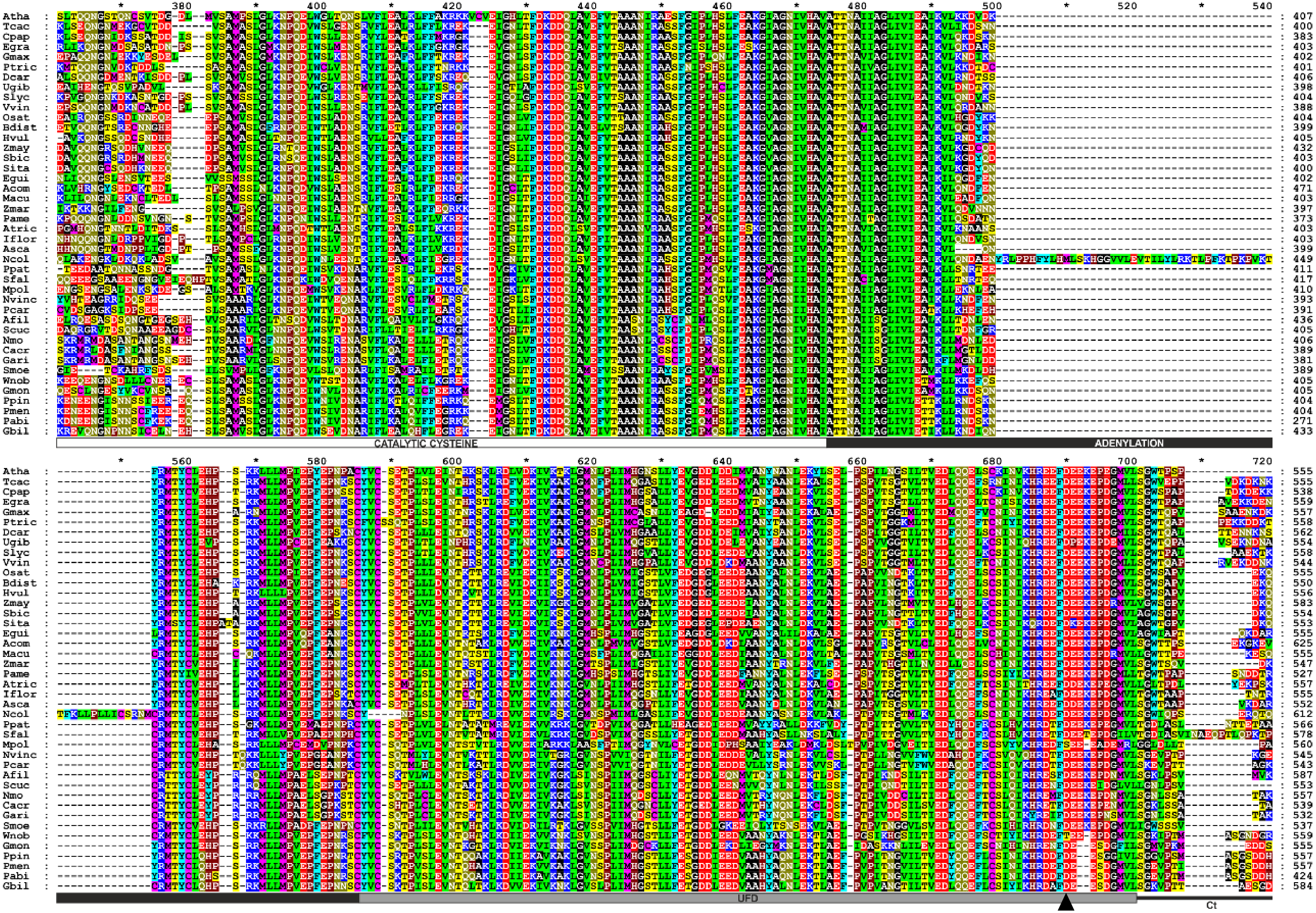
Atha
Tcac
Cpap
Egra
Gmax
Ptric
Dcar
Ugib
Slyc
Vvin
Osat
Bdist
Hvul
Zmay
Sbic
Sita
Egui
Acom
Macu
Zmar
Pame
Atric
Iflor
Asca
Ncol
Ppat
Sfal
Mpol
Nvinc
Pcar
Afil
Scuc
Nmo
Cacr
Gari
Smoe
Wnob
Gmon
Ppin
Pmen
Pabi
Gbil
CATALYTIC CYSTEINE
ADENYLATION
UFD
Ct

04

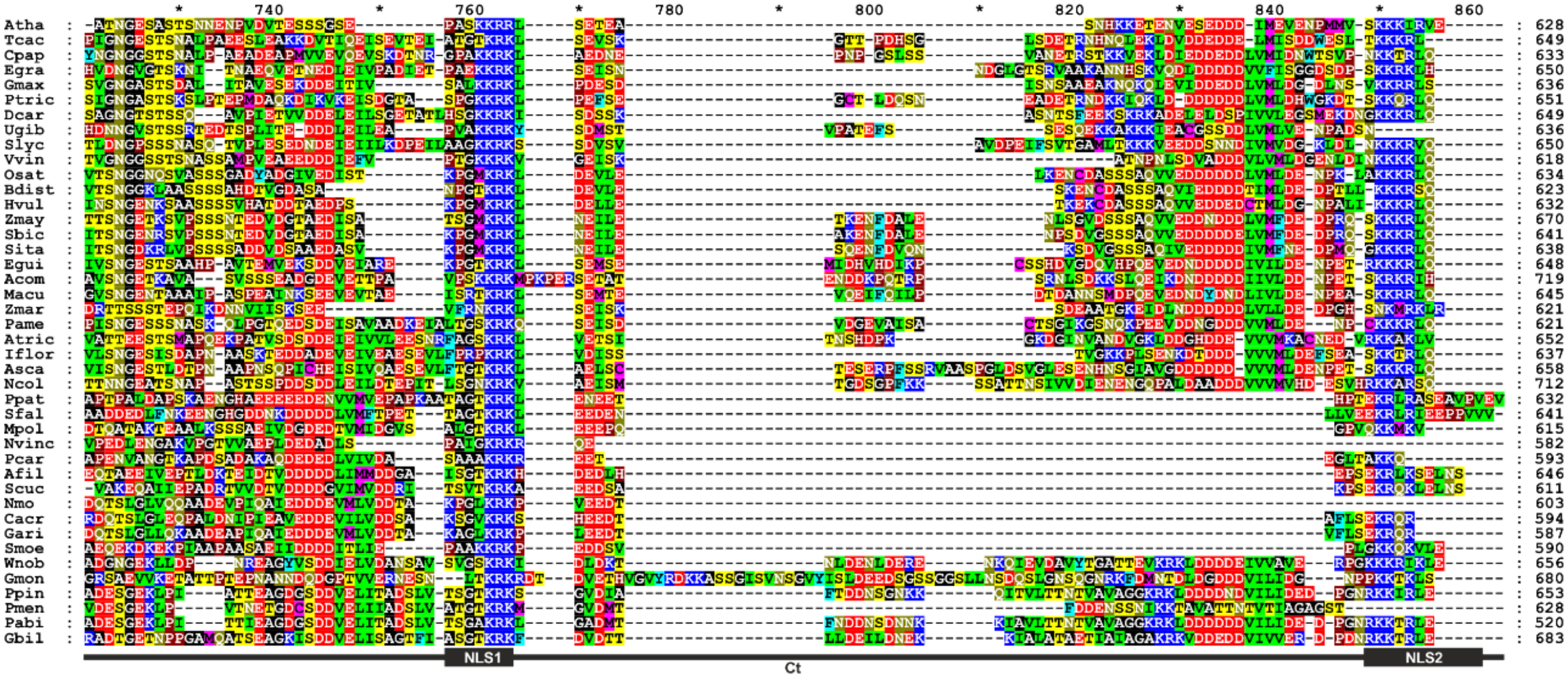
Atha
Tcac
Cpap
Egra
Gmax
Ptric
Dcar
Ugib
Slyc
Vvin
Osat
Bdist
Hvul
Zmay
Sbic
Sita
Egui
Acom
Macu
Zmar
Pame
Atric
Iflor
Asca
Ncol
Ppat
Sfal
Mpol
Nvinc
Pcar
Afil
Scuc
Nmo
Cacr
Gari
Smoe
Wnob
Gmon
Ppin
Pmen
Pabi
Gbil
NLS1
Ct
NLS2

**Supplementary Figure 9: Multiple alignment of SAE2 protein sequences from 42 land plant species.** SAE2 functional domains are indicated by boxes below the alignment and their positions are defined in reference to the *Arabidopsis* sequence. The arrow indicates the catalytic cysteine and the arrow head indicates the position of the first amino acid encoded by the last exon. Alignment is colored according to amino acid identities in all sequences. Atha, *Arabidopsis thaliana*; Tcac, *Theobroma cacao*; Cpap, *Carica papaya*; Egra, *Eucalyptus grandis*; Gmax, *Glycine max*; Ptri, *Populus trichocarpa*; Dcar, *Daucus carota*; Ugib, *Utricularia gibba*; Slyc, *Solanum lycopersicum*; Vvin, *Vitis vinifera*; Osat, *Oryza sativa ssp. japonica*; Bdis, *Brachypodium distachyon*; Hvul, *Hordeum vulgare*; Zmay, *Zea mays B104*; Sbic, *Sorghum bicolor*; Sita, *Setaria italica*; Egui, *Elaeis guineensis*; Acom, *Ananas comosus*; Macu, *Musa acuminata*; Macu, *Musa acuminata*; Zmar, *Zostera marina*; Pame, *Persea americana Hass*; Atri, *Amborella trichopoda*; Iflo, *Illicium_floridanum*; Asca, *Austrobaileya_scandens*; Ncol, *Nymphaea colorata*; Wnob, *Wollemia nobilis*; Gmon, *Gnetum montanum*; Ppin, *Pinus pinaster*; Pmen, *Pseudotsuga menziesii*; Pmen, *Pseudotsuga menziesii*; Pabi, *Picea abies*; Gbil, *Ginkgo biloba*; Afil, *Azolla filiculoides*; Scuc, *Salvinia cucullata*; Nmon, *Notholaena_montieliae*; Cacr, *Cryptogramma_acrostichoides*; Gari, *Gaga_arizonica*; Nvin, *Nothoceros vincentianus*; Pcar, *Phaeoceros_carolinianus*; Smoe, *Selaginella moellendorffi*; Ppat, *Physcomitrella patens*; Sfal, *Sphagnum fallax*; Mpol, *Marchantia polymorpha*.
